# HIV infection does not alter Interferon α/β receptor expression on mucosal immune cells

**DOI:** 10.1101/688440

**Authors:** Julia Ickler, Sandra Francois, Marek Widera, Mario L. Santiago, Ulf Dittmer, Kathrin Sutter

## Abstract

The innate immune response induced by type I interferons (IFNs) play a critical role in the establishment of HIV infection. IFNs are induced early in HIV infection and trigger an antiviral defense program by signaling through the IFNa/b receptor (IFNAR), which consists of two subunits, IFNAR1 and IFNAR2. Changes in IFNAR expression in HIV target cells, as well as other immune cells, could therefore have important consequences for initial HIV spread. It was previously reported that IFNAR2 expression is increased in peripheral blood CD4^+^ CXCR4^+^ T cells of HIV^+^ patients compared to HIV uninfected controls, suggesting that HIV infection may alter the IFN responsiveness of target cells. However, the earliest immune cells affected by HIV *in vivo* reside in the gut-associated lymphoid tissue (GALT). To date, it remains unknown if IFNAR expression is altered in GALT immune cells in the context of HIV infection and exposure to IFNs, including the 12 IFNa subtypes. Here, we analyzed the expression of surface bound and soluble IFNAR2 on Lamina propria mononuclear cells (LPMCs) isolated from the GALT of HIV^−^ individuals and in plasma samples of HIV^+^ patients. IFNAR2 expression varied between different T cells, B cells and natural killer cells, but was not altered following HIV infection. Furthermore, expression of the soluble IFNAR2a isoform was not changed in HIV^+^ patients compared to healthy donors, nor in LPMCs after HIV-1 infection *ex vivo*. Even though the 12 human IFNα subtypes trigger different biological responses and vary in their affinity to both receptor subunits, stimulation of LPMCs with different recombinant IFNα subtypes did not result in any significant changes in IFNAR2 surface expression. Our data suggests that potential changes in the IFN responsiveness of mucosal immune cells during HIV infection is unlikely dictated by changes in IFNAR2 expression.

## Introduction

Natural transmission of HIV occurs via the vaginal or the gastrointestinal mucosa, where first responses against invading virus are mounted. Following HIV infection, the gastrointestinal mucosa succumbs to profound enteropathy, including increased gastrointestinal inflammation, malabsorption, diarrhea and increased intestinal permeability [1], as well as microbial translocation and dysbiosis [2,3]. The gastrointestinal lymphoid tissue (GALT) shows the strongest substantial depletion of CD4^+^ T cells, occurring mostly in CD4^+^ CCR5^+^ T cells [4,5] in comparison to peripheral mononuclear blood cells (PBMCs) which are commonly used for studying HIV immune responses. An earlier study conducted with simian immunodeficiency virus (SIV) proved the CD4^+^ T cell depletion not only to be more severe, but also to occur much faster in the GALT [6] and showed a delayed restoration of CD4^+^ T cell numbers following early combined antiretroviral therapy (cART) initiation in comparison to PBMCs [7]. As Lamina propria mononuclear cells (LPMCs) can be infected efficiently by CCR5-tropic HIV-1 strains without the need for exogenous stimulation, the Lamina propria aggregate culture (LPAC) can be used as an efficient *ex vivo* model to study HIV infection close to the physiological background [8].

Since adaptive immunity has yet to be mounted in the early stages of infection, innate immune responses are of great importance as the first line of defense. Early host immune responses in the GALT are mediated by type I interferons (IFNs), which are mainly secreted by plasmacytoid dendritic cells (pDC) [9]. Type I IFNs are a pleiotropic cytokine family consisting of IFNα, IFNβ, IFNε, IFNκ and IFNω. The human chromosome 9 contains 13 genes encoding for 12 individual IFNα subtypes [10], highly conserved proteins with an amino acid sequence homology of 75-99 % [11]. All type I IFNs bind to the common IFNα/β receptor (IFNAR), which is widely expressed on most cell types [12]. The receptor consists of two subunits, IFNAR1 and IFNAR2, which associate with Janus kinases (Jak) Tyk2 (IFNAR1) and Jak1 (IFNAR2). Upon initial ligand binding by IFNAR2, IFNAR1 is recruited and subsequent to formation of the ternary complex out of IFNAR1, IFNAR2 and IFNα or IFNβ, Tyk2 and JAK1 become activated. Type I IFN signal transduction commonly takes place via the classical Jak-STAT pathway leading to the transcription of numerous IFN-stimulated genes (ISGs), among them several genes that encode for proteins with direct antiviral activity, e.g., apolipoprotein B mRNA editing enzyme, catalytic polypeptide like (APOBEC), SAM domain and HD domain containing protein 1 (SamHD1) and interferon induced GTP binding protein MX2 (MX2). Additionally, IFNα subtypes indirectly exert immunomodulatory functions and anti-proliferative effects on cells of the innate and adaptive immune system [13]. The IFNAR2 subunit is of special interest, as it is responsible for initial ligand binding and three different isoforms are described. IFNAR2c contains long intracellular domains with associated kinases and is responsible for signal transduction. IFNAR2b is likewise a membrane bound isoform lacking the intracellular domains, which is thought to be a negative regulator for type I IFN signaling [14]. Meanwhile, IFNAR2a is a soluble isoform, produced by either alternative splicing or by proteolytic cleavage at the cell surface and, in the murine organism, independently regulated from trans-membranous IFNAR2 [15]. It is known to be increased in multiple sclerosis[16], adeno carcinoma and lung cancer [17,18] and was found to be elevated and negatively correlated with successful IFN therapy in Hepatitis C patients [19].

Compared to healthy individuals, IFNα expression is increased and the specific expression pattern of IFNα subtypes is changed in HIV^+^ patients. [20–22]. Even though the exact role of IFNs in HIV infection is still under debate, beneficial implications of IFNα as suppressed viral load, increased NK cell function and enhanced suppressive capacity of CD8^+^ T cells as well as an increase in the expression of ISGs containing HIV restriction factors and thus hindering viral transmission and replication could be observed [23–25]. On the other hand, rescued CD4^+^ T cell depletion and restored cell function following blockade of IFNAR was reported [23,26], as well as systemic immune activation and limited antigen-specific T cell responses [24], thus indicating a potential detrimental influence of IFN on the course of HIV infection.

Due to their high antiviral potential, several studies testing IFNα as a treatment option for HIV were conducted, but patients showed no or only mild benefits from the treatment [27–29]. This, along with the development of highly effective antiretroviral therapies, led to decreased interest in IFNα as a potential therapeutic strategy against HIV [30]. Furthermore, several studies showed that the clinically approved IFNα2 subtype possesses only weak antiviral activity against HIV [9,31]. Thus the question remains, wether an IFNα subtype with higher antiviral capacities against HIV, such as IFNα14, might be of more use against HIV, for example as a potential addition to cART. Initial studies with humanized mice using IFNα14 as an antiviral agent showed promising results. In contrast to IFNα2 treatment, IFNα14 was able to reduce viremia and proviral loads in post-exposure prophylaxis and treatment of acute infection and reduced hyperimmune activation [31]. Furthermore, gene therapy with plasmids encoding for IFNα14, but not for IFNα2, was shown to provide long-term suppression of HIV replication [32].

Even though the individual IFNα subtypes exert different biological responses, they all bind to the same receptor. One factor possibly influencing the diverse biological outcome is the different affinity of IFNα subtypes to both receptor subunits IFNAR1 and IFNAR2 [33]. For IFNAR2, a correlation between antiviral activity and receptor affinity was observed [9], while anti-proliferative effects seem to be more associated with the affinity to IFNAR1 and the stability of the ternary complex, respectively [33,34].

Various factors regulate the biological response to IFNα stimulation. The affinity of the individual IFNα subtype to both receptor subunits und their cell surface density is pivotal. Combined with the dose of IFNα the cell is receiving, the overall avidity of an individual IFNα subtype is a major determinant for the biological outcome [35,36]. Cell-type specific effects as well as microenvironment based factors, e.g. downregulation of IFNAR1 in Influenza A infections [37] or in colorectal cancer [38] also modulate the IFNα response. Additionally, the precise timing of exposure to IFNα in reference to proceeding or subsequent priming of the target cells as well as the duration of ligand binding shape the biological outcome [35]. Ligand-binding induced receptor downregulation is described for both receptor subunits on different cell lines following stimulation with IFNα2 and IFNβ [39]. According to its higher affinity to the receptor, IFNβ stimulation decreased the receptor surface expression to much stronger extents [39].

Yoder *et al.* showed reduced expression of canonical ISGs in LPMCs, infected with HIV [40], which might indicate differences in the IFN responsiveness following HIV infection. Another study observed differences in the ISG expression pattern following stimulation with different IFNα subtypes [9]. Changes in IFNAR expression during infection or subsequent to stimulation with different IFNα subtypes would have the potential to modify IFN responsiveness and induce differences in the ISG expression pattern. A recent study by Killian *et al*. observed increased IFNAR2 expression on CXCR4^+^ CD4^+^ T cells in peripheral blood of HIV^+^ patients. The first cells to encounter HIV are CCR5 CD4^+^ T cells in the GALT but so far, there has been no study to compare IFNAR2 expression on CCR5^+^ CD4^+^ T cells in LPMCs from PLWH and healthy donors. Up to date it is also unknown, if IFNα subtypes shape the IFNα response by differences in their potential to regulate receptor expression. To mirror the physiological background of HIV infection, we performed experiments using LPMCs isolated from the GALT of otherwise healthy patients, undergoing abdominal surgery. Here, we determine the IFNAR2 expression on different immune cell subsets and investigate the influence of HIV infection on the expression of IFNAR2a and IFNAR2b/c. Varying levels of IFNAR2b/c expression on different immune cell subsets were observed, with the highest expression occurring in B cells, followed by NK and CD4^+^ and CD8^+^ T cells. HIV infection did not change the expression of either the surface bound or the soluble isoform of IFNAR2. Finally, stimulation with a half maximal effective concentration (EC_50_) of different IFNα subtypes did not lead to any changes in receptor surface expression.

## Material & Methods

### LPMCs

LPMCs were isolated from patients undergoing abdominal surgery at the University Hospital Essen. Samples were received via the Westdeutsche Biobank, pathologically evaluated and judged macroscopically normal. Gut samples from 16 different patients where used in this study, 6 of them female, 10 male. The average age was 55.8 years (+/-13.2 years). In order to isolate LPMCs, samples were processed as previously described [41]. Cells were cultivated in RPMI1640 with 10% human serum (Type AB, Pan Biotech, Aidenbach, Germany), 1% Penicillin/Streptomycin (Thermo Fisher, Waltham, USA), 1% L-Glutamine (Thermo Fisher, Waltham, USA) and 0.4% Piperacillin/Tazobactam (Fresenius Kabi, Bad Homburg an der Höhe, Germany). Cells were incubated at 1×10^6^ cells/ml at 37 °C and 5% CO_2_ for the indicated time period. LPMC collection was approved by the Ethics Committee (No.:15-6310) of the medical faculty at the University of Duisburg-Essen.

### Infection

Virus stocks were produced and quantified as described elsewhere [42]. Infection experiments were performed using R5-tropic HIV-1_NL4.3_. LPMCs were thawed one day prior to infection. Cells were infected with an MOI of 0.005, 0.01 or 0.02 via spinoculation at 1000 g and cultivated with 1×10^6^ cells/ml. Supernatant of HIV-infected and mock-infected cells were collected at the indicated time points and stored at −80°C until further use.

### TZM-bl Assay

TZM-bl cells (7×10^3^ cells/100 µl) were plated in a 96 well plate. 24 h later, the media was removed and serial dilutions of virus stock or 100 µl supernatant of cultivated LPMCs were added to the cells. Two days post infection, supernatant was removed, the cells were washed with cold PBS and subsequently fixed with 0,88% formaldehyde and 0,0625% glutaraldehyde in PBS. The cells were again washed with 200 µl cold PBS and 100 µl staining solution per well (400 mM K3[Fe(CN)6], 400 mM K4[Fe(CN)6], 400 mM MgCl, 20 mg/ml X-Gal) was added. 24 h later, the assay was analyzed by counting wells that contained blue stained cells.

### p24-ELISA

The p24 ELISA with supernatant from infected and non-infected LPMC cultures was performed according to the manufacturer’s protocol (R&D Systems, Minneapolis, USA). Absorbance was measured using the Spark^®^ 10M multimode microplate reader (Tecan, Männedorf, Switzerland).

### FACS staining

Cell surface staining was performed using the following antibodies: anti-CD3 (SK7, BioLegend), anti-CD8 (RPA-T8, BioLegend, San Diego, USA), anti-CD19 (SJ25C1, BioLegend), anti-CD56 (NCAM16.2, BioLegend), anti-IFNAR2 (REA124, Miltenyi Biotech GmbH, Bergisch Gladbach, Germany), Fc Block human (Miltenyi Biotech GmbH). For surface staining, cells were washed once with FACS Buffer (PBS containing 0,1% bovine serum albumin and 0,02% sodium azide), Pellets were then resuspended and incubated for 10 min with antibodies mixture. Cells were washed once more with FACS Buffer and stored until acquisition. Dead cells were excluded from the analysis using Zombie Dye (BioLegend). Samples were acquired with a BD LSR II flow cytometer and a BD FACS Canto II flow cytometer (Becton Dickinson, Franklin Lakes, USA) and data were analysed using FACSDiva and FlowJo Version 10 (both Becton Dickinson).

### sIFNAR2a ELISA

Plasma samples of HIV^+^ patients (22 cART-naïve as well as 22 cART-experienced HIV^+^ patients) were collected from the clinics of dermatology at the University Hospital Essen; HIV^−^ samples (n=13) were donated by healthy individuals of the University Hospital Essen. Blood collection was approved by the Ethics Committee (No.:11-4715) of the medical faculty at the University of Duisburg-Essen. Viral loads among cART naïve patients ranged from 2768 copies/ml to 460,400 copies/ml with a mean of 162,643 copies/ml, +/-155,875 copies/ ml. Supernatants of HIV-infected LPMCs were sampled as indicated above. The ELISA for sIFNAR2a was performed according to the manufacturer’s protocol, the detection limit was 0.16 ng sIFNAR2a (RayBiotech, Norcross, USA). Absorbance was measured using the Spark^®^ 10M multimode microplate reader (Tecan, Männedorf, Switzerland).

### Stimulation with different human IFNα subtypes

IFNα subtypes were produced and purified as previously described [31]. The activity of each subtype was determined using the human ISRE-Luc reporter cell line, a retinal pigment epithelial cell lines transfected with a plasmid containing the Firefly Luciferase gene stabily integrated under control of the IFN-stimulation-response element (ISRE). Following stimulation with IFNα, chemiluminescence can be detected and used to calculate the respective activity against commercially available IFNα (PBL assays sciences, Piscataway, USA) [31]. For stimulation experiments, thawed LPCMs were cultivated for the indicated time points with the EC_50_ of each subtype (see Fig. 3): IFNα1 1.9×10^3^ U/ml, IFNα2 1×10^3^ U/ml, IFNα8 4×10^3^ U/ml, IFNα14 6.5×10^1^ U/ml.

### Statistical Analysis

Statistical analysis and graphical presentations were carried out with Graph Pad Prism v6. For comparisons of two datasets, paired t-test was performed. For data including several groups one way ANOVA and Bonferroni’s multiple comparison test was performed.

## Results

### IFNAR2 expression after HIV-1 infection

The gut-associated lymphoid tissue is a major site of HIV replication and spread shortly after transmission due to the high numbers of activated, effector memory CD4^+^ T cells that express the co-receptor CCR5 in this compartment. This process highly depends on type I IFN mediated innate immune responses. The IFN α/β receptor consists of the two subunits IFNAR1 and IFNAR2, of which IFNAR2 is responsible for initial ligand binding. To analyze the impact of HIV-1 infection on IFNα signaling, we determined the surface expression of IFNAR2 on NK cells, B cells, CD4^+^ and CD8^+^ T cells isolated from the gastrointestinal mucosa of 11 healthy donors, undergoing abdominal surgery. All gut samples were pathologically evaluated and judged macroscopically normal by a pathologist. The percentages of IFNAR2-expressing cells varied strongly between different immune cell subsets, with B cells containing almost three times more IFNAR2-expressing cells (mean 54.7 %) than NK cells (mean 20.3 %). The smallest proportions of IFNAR2^+^ cells was found among CD4^+^ T cells (mean 12.3 %) and CD8^+^ T cells (mean 13.8 %) (Fig.1). Analyzing the mean fluorescence intensity (MFI) at a single cell level, few differences between immune cell subsets were observed with individual NK cells expressing only slightly higher IFNAR2 levels, followed by B cells and both CD4^+^ T cells and CD8^+^ T cells. Successful infection of LPMCs was examined with TZM-bl assay and p24 ELISA. Following infection with HIV-1_NL4.3_ for four or seven days (data not shown) at different MOIs (MOI 0.005, MOI 0.01 and MOI 0.02), neither the percentages of IFNAR2 expressing cells between immune cell subsets, nor the amount of IFNAR2 molecules at the single cell level (MFI, data not shown) changed, indication that HIV itself has no effect on IFNAR2 surface expression.

**Fig. 1.**
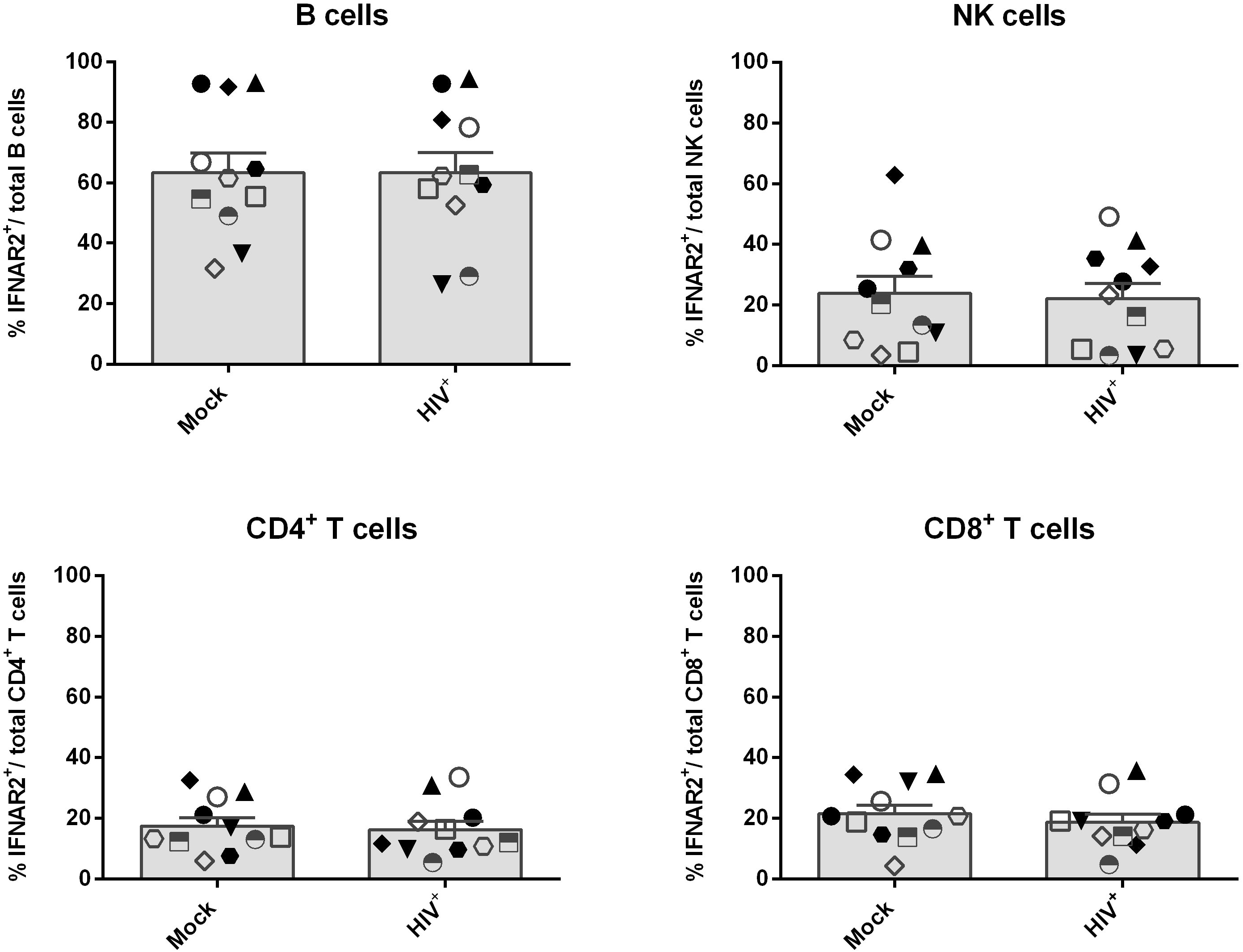
IFNAR2 expression following HIV-1 infection of LPMCs. LPMCs from healthy donors were infected with HIV-1_NL4.3_ and cultivated for 4 days. Surface expression of IFNAR2 on mock treated and infected cells was determined via flow cytometry on CD4^+^ T cells, CD8^+^ T cells, B cells and NK cells. Individual frequencies of IFNAR2-expressing cells and mean values (+SEM) are shown as dots and bars (n=11). Differences between the groups were analyzed by paired student’s t test.

### Expression of IFNAR2 isoforms

The IFNAR2 receptor subunit is expressed in three different isoforms: IFNAR2a, IFNAR2b and IFNAR2c. IFNAR2c and IFNAR2b are membrane-bound isoforms, IFNAR2c being responsible for signal transduction while IFNAR2b, which lacks the necessary intracellular domains, is thought to be a negative regulator for type I IFN signaling [14]. IFNAR2a is a soluble isoform, which can be produced through alternative splicing or proteolytic cleavage from the cell surface. Since we could not observe any difference in IFNAR2b/c surface expression following HIV-infection (Fig. 1), we next analyzed the expression of soluble IFNAR2 (sIFNAR2a) in the supernatant of mock-treated and HIV-infected LPMCs at days 1, 3, 4, 5, 6 and 7 post infection via ELISA (Fig. 2a, data shown for day 4). Similar to the surface expression of IFNAR2 (Fig. 1), we did not observe any significant differences in the sIFNAR2a expression between mock-treated (mean 0.3 ng/ml) and HIV-infected LPMCs (mean 0 ng/ml) (Fig. 2a). To examine possible effects of HIV-1 infection *in vivo*, we compared the sIFNAR2 expression in plasma samples of healthy donors (mean 0.41 ng/ml) with plasma samples of HIV-1^+^ cART-naïve (mean 0.40 ng/ml) and cART-experienced patients (mean 0.21 ng/ml) (Fig. 2b). According to the *in vitro* HIV-infected LPMC results, sIFNAR2a levels did not significantly differ between the groups. Furthermore, we did not find any correlation between viral loads and sIFNAR2a expression in cART-naïve patients (data not shown). Additionally, no association between the cART regimen and sIFNAR2a expression was observed in cART-experienced HIV^+^-patients (data not shown).

**Fig. 2.**
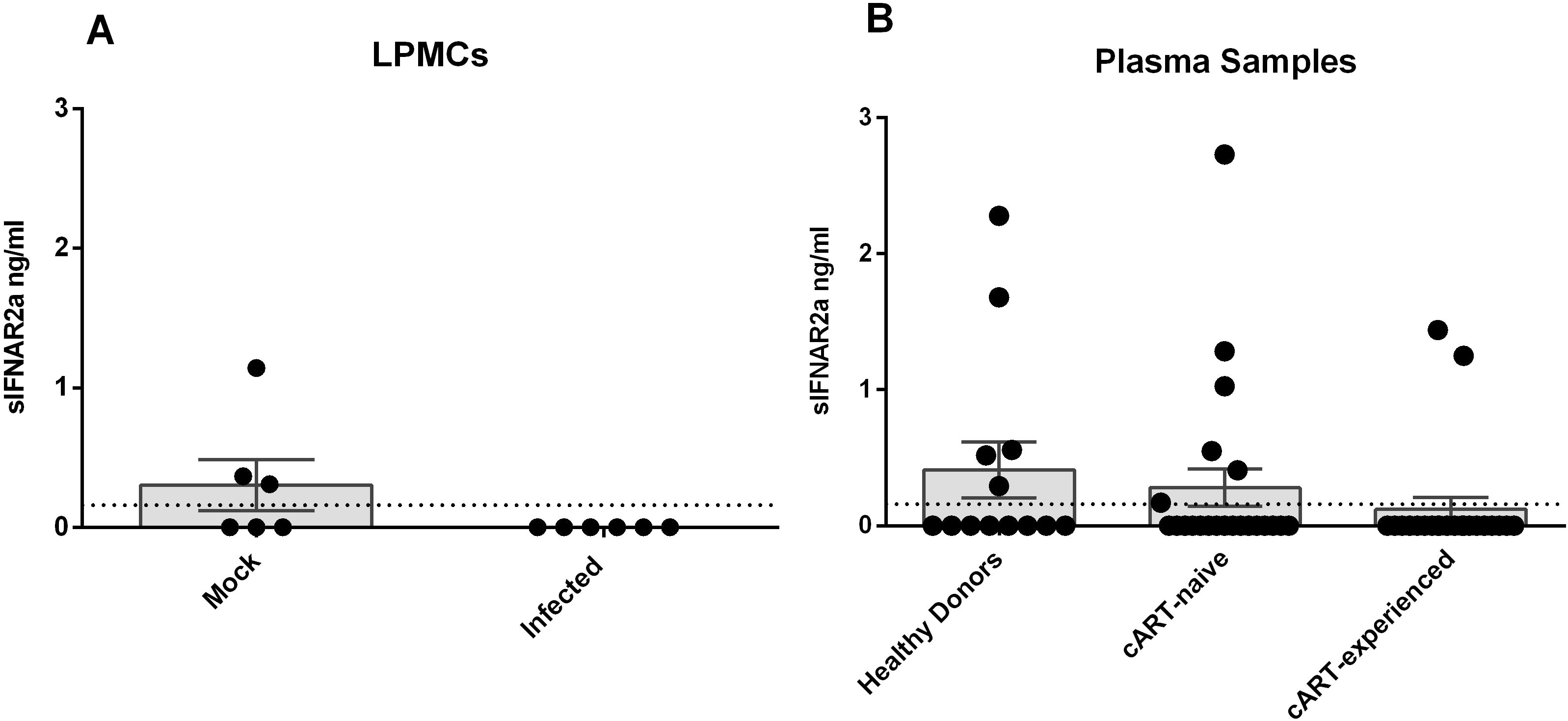
Soluble IFNAR2a expression in HIV-1 infected LPMCs and plasma samples. Supernatant of mock-treated and HIV-infected LPMCs from 4 days post infection (A, n=6) and plasma samples (B) from healthy donors (n=14), cART-naïve (n=22) and cART-experienced HIV-1^+^ patients (n=22) were analyzed for sIFNAR2a expression via ELISA. The dotted line indicates the detection limit of 0.16 ng/ml. Individual frequencies of sIFNAR2a-expressing samples and mean values (+SEM) are shown as dots and bars. Statistically significant differences between the groups were analyzed by student‘s t test (LPMCs) and ordinary one way ANOVA analysis with Bonferroni’s multiple comparison (plasma samples).

### IFNAR2 expression following IFNα subtype stimulation

IFNAR1 and IFNAR2 are known to be differentially regulated and their expression can decrease upon ligand binding [39]. Since the 12 individual human IFNα subtypes differ in their antiviral capacity against HIV [9,31], we sought to compare subtypes with very high (IFNα8 & IFNα14) to those with very low (IFNα1 & IFNα2) antiviral activities, with respect to their potential to influence receptor expression. Furthermore, we chose IFNα subtypes representing subtypes high with (α14, α2), middle (α8) and low (α1) affinity to IFNAR2.

LPMCs from four different healthy donors were stimulated with the half maximally effective concentration (EC_50_; Fig. 3) of IFNα1, IFNα2, IFNα8 and IFNα14 for 15 min, 30 min, 2 h and 24 h. Subsequently, surface receptor expression was determined via flow cytometry. We did not see significant differences in the surface expression of IFNAR2 on CD4^+^ and CD8^+^ T cells, B cells and NK cells after stimulation with the different IFNα subtypes. Furthermore, stimulation did not lead to changes in IFNAR2 expression compared to the unstimulated control. Slight elevations of IFNAR2 expression up to approximately 130% compared to the unstimulated control were observed 24 h following stimulation for all tested cell populations. Interestingly, one of the four tested LPMC donors exhibited an increase of IFNAR2 expression of 270% on CD8^+^ T cells when stimulated (Fig. 4). In addition to the percentages of IFNAR2 expressing cells, no differences were detected at the single cell level when analyzing the MFI (data not shown).

**Fig. 3.**
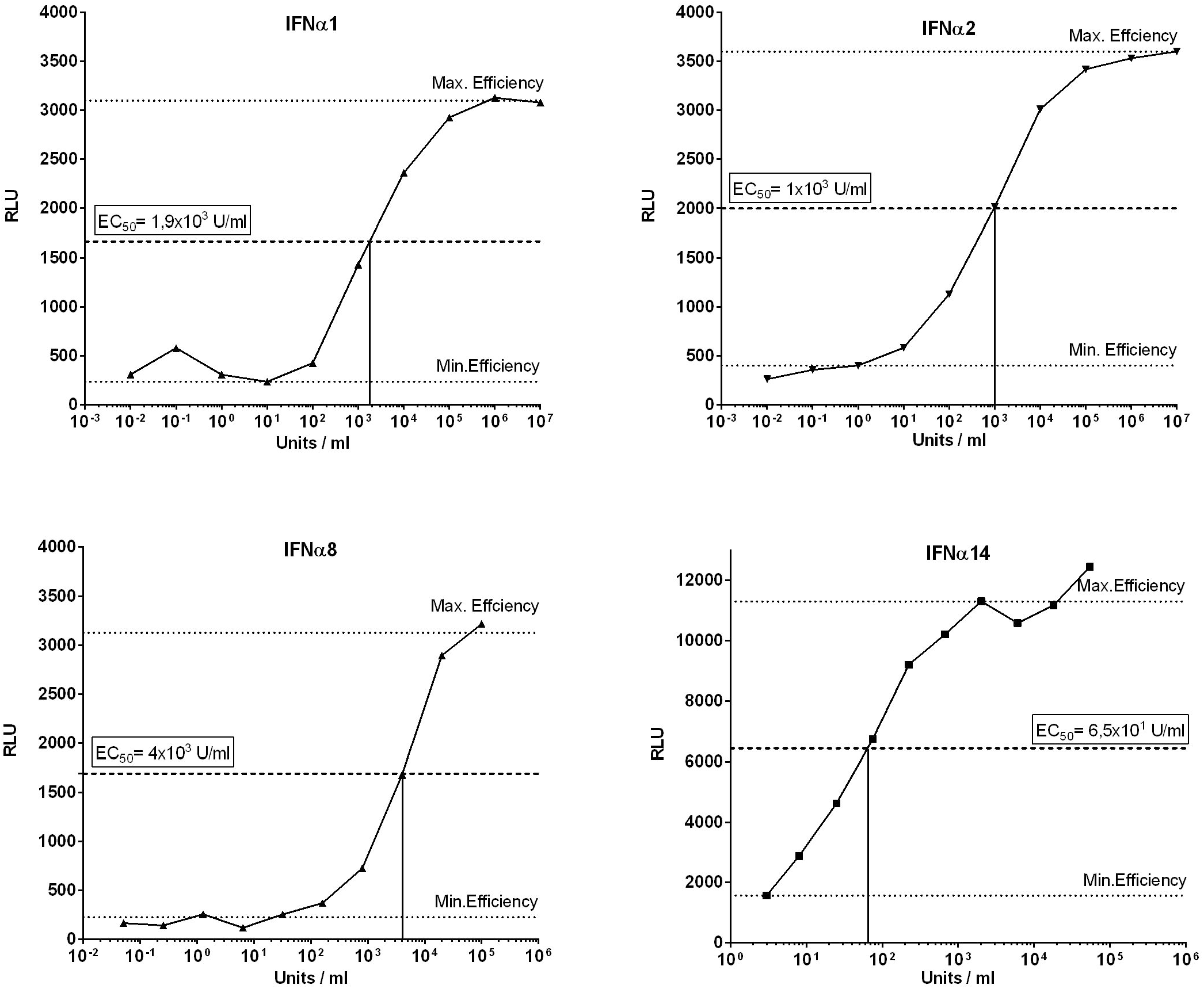
Titration of individual IFNα subtype activity. The recombinant human reporter cell line ISRE-Luc was stimulated with series dilutions of each IFNα subtype. Maximal and minimal effective concentrations were determined and the half maximal effectivity concentration (EC_50_) was calculated. The upper and lower dotted lines indicate maximal and minimal effective concentration of the respective IFNα subtype, the middle dotted line indicates the EC_50_.

**Fig. 4.**
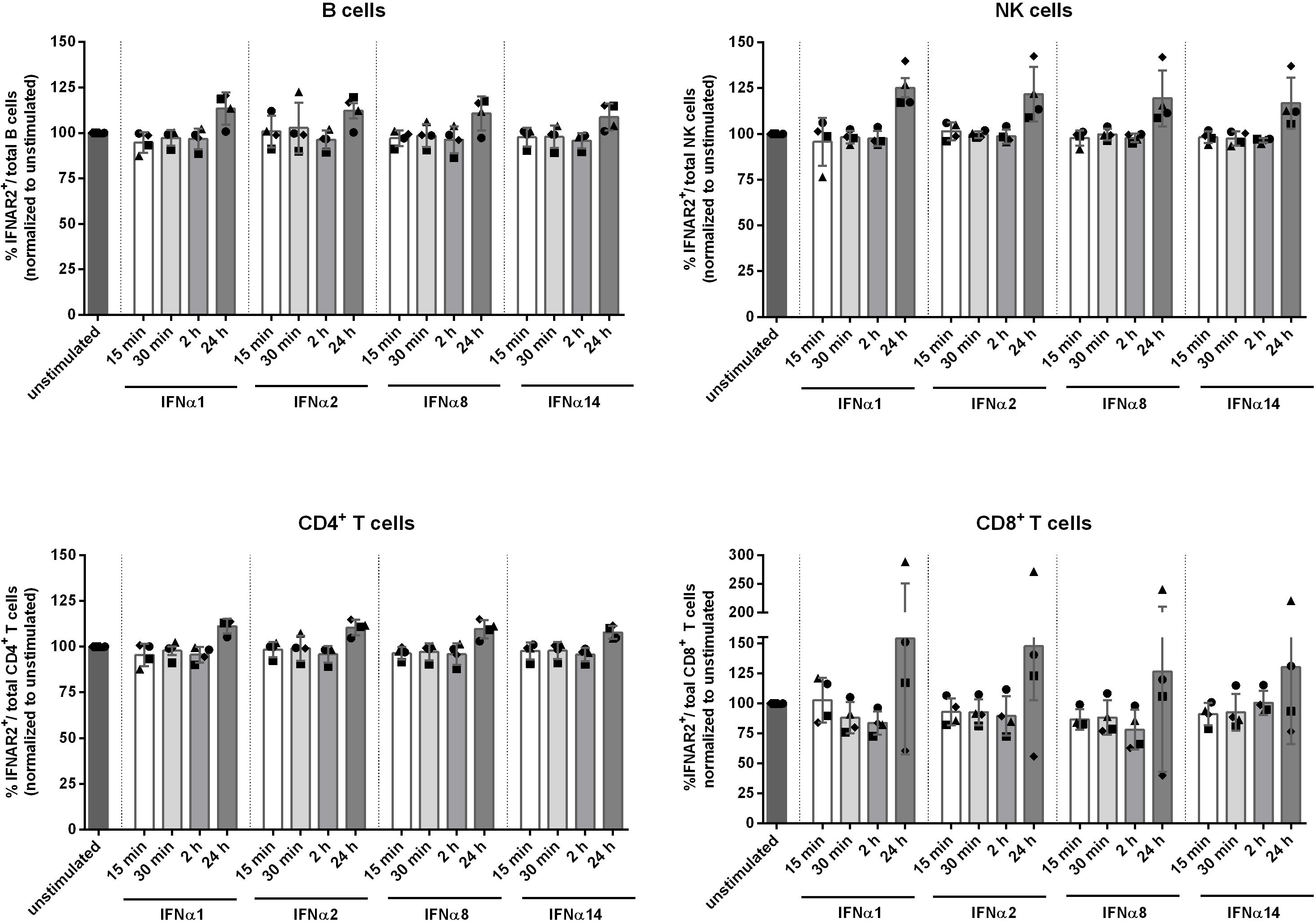
IFNAR2 expression following stimulation with IFNα subtypes. LPMCs were stimulated with the EC_50_ of IFNα1, IFNα2, IFNα8 and IFNα14 for 15 min, 30 min, 2 h and 24 h. Surface expression of IFNAR2 on unstimulated and stimulated cells was determined via flow cytometry on CD4^+^ T cells, CD8^+^ T cells, B cells and NK cells. Individual frequencies of IFNAR2-expressing cells and mean values (+SEM) are shown as dots and bars (n=4). Differences between the groups were analyzed by ordinary one way ANOVA analysis and Bonferroni’s multiple comparisons.

## Discussion

IFNα/β responses are the immediate early host response against invading viruses and crucial for mounting immune responses. Upon engaging IFNAR, signal transduction results in the expression of hundreds of IFN-stimulated genes, among them several genes with direct antiviral functions. Even though all 12 human IFNα subtypes bind the same receptor, they trigger different biological responses and vary strongly in their antiviral activity against HIV. Whereas the only clinically approved subtype IFNα2 exhibits weak antiviral effects, other subtypes, such as IFNα14, are highly antiviral against HIV [9,31]. In this study, we aimed to determine whether the different responses are based on differential expression of IFNAR2 on various immune cell populations and if receptor expression levels might be altered upon infection with HIV. Indeed we could find strong differences in the percentages of IFNAR2 expressing cells, which were almost three times as high on B cells compared to CD4^+^ and CD8^+^ T cells (Fig. 1). These findings are in agreement with other studies, which also showed higher expression of IFNAR2 on B and NK compared to T cells [43,44]. Whereas Killian *et al.* [45] observed increased IFNAR2 expression on CXCR4^+^ CD4^+^ T cells in whole blood of HIV^+^ patients compared to healthy donors, we did not see any differences in IFNAR2 surface expression on the analyzed immune cell subsets following HIV infection of LPMCs *in vitro*. Taken into account that the biological effect of IFNα subtypes highly depends on the microenvironment and cell type [35], differences in receptor expression between different tissues are not unexpected. Furthermore, Killian *et al*. found increased IFNAR2 expression only on CXCR4^+^ CD4^+^ T cells of HIV^+^ patients compared to healthy donors, but not on CCR5^+^ CD4^+^ T cells. We did not determine the ratio of CXCR4^+^ CD4^+^ T cells versus CCR5^+^ CD4^+^ T cells in our samples. However, while only a small proportion of peripheral CD4^+^ T cells express CCR5, the majority of CD4^+^ T cells in LPMCs are reported to express CCR5 [8]. For HIV^+^ patients, increased IFNAR2 expression was associated with clinical progression despite effective antiviral therapy progression [45]. Perhaps additional time points after infection are necessary to detect significant changes in IFNAR2 expression when using the LPAC model. Finally, Killian *et al.* only saw a significant increase in IFNAR2 expression on HAART-patients, and not on therapy-naïve long-term non-progressors, possibly indicating that HAART have an influence on IFNAR2 expression rather than HIV itself [45]. Other studies in mouse models reported a downregulation of IFNAR1 and IFNAR2 during influenza A infections, caused by viral proteins [37,46]. Influenza A was also shown to downregulate IFNAR1 and IFNAR2 in human monocyte-derived macrophages, however *ex vivo* infection of human lung tissue with Influenza A only reduced the expression of IFNAR1, but not IFNAR2 [46]. Among the two subunits, IFNAR1 expression seems to be more stringently regulated [12] and stable expression on the cell surface requires the binding of Tyk2 to its intracellular domains [47,48]. While IFNAR1 is directed towards lysosomal degradation following downregulation from the cell surface, the fate of IFNAR2 seems to depend on the ligand: Following IFNα stimulation, IFNAR2 is rapidly recycled back to the cell surface, though it is targeted for degradation upon binding of IFNβ [49]. By comparing IFNAR1^−/−^ and IFNAR2 ^−/−^ mouse models, distinct roles for each subunit in regulating the immune response against Influenza A could be discerned. Whereas the full receptor was needed for complete protection against Influenza A infection, higher morbidity, mortality and increased viral burden in IFNAR2^−^/^−^ mice indicate a more extensive role of IFNAR2 in antiviral immunity. Furthermore, stimulation with IFNβ was able to induce protection from lethal Influenza A infection in IFNAR1^−^/^−^ mice, but not in IFNAR2^−^/^−^ mice. In comparison, IFNAR2^−^/^−^ mice were better able to control bacterial superinfections subsequent to Influenza A infection [50]. FNβ is reported to be able to bind IFNAR1 independently from IFNAR2 [51], which further strengthens the possibility of distinct roles for reach receptor subunit. Also in HIV, different functions and regulation for IFNAR1 and IFNAR2 are possible. As the majority of data concerning IFNAR1 and IFNAR2 expression and regulation were generated using mouse models or cell lines, one has to be careful with translating these findings to humans. Though IFNα/β signaling plays an absolutely crucial role in mounting immune responses and protection against viral infection in mice, a recent translational report from Duncan and colleagues about a patient deficient in IFNAR2 challenged this perspective, stating that as long as infections in humans are local and not systemic, lack of IFNα/β signaling has no effect on the development of T cells and does not increase the susceptibility towards respiratory infections [52].

Our findings concerning the expression of the soluble isoform of IFNAR2, sIFNAR2a, are in accordance with the results for IFNAR2b/c surface expression. We detected trace amounts of sIFNAR2a in the supernatant of HIV-infected LPMCs (Fig. 2A) and we did not see any difference in sIFNAR2a concentrations in the plasma of healthy donors and HIV^+^ cART-naïve and cART-experienced patients (Fig. 2B).

Both agonistic and antagonistic properties of sIFNAR2a where shown in mice, specifically their ability to influence the efficacy of therapeutic IFNα treatment [15,53]. In mouse cell lines and primary cells overexpressing sIFNAR2a, the antiproliferative and antiviral effects of IFNα and IFNβ stimulation were inhibited. Recombinant sIFNAR2a was able to bind IFNα and IFNβ and complex with IFNAR1 on the surface of IFNAR2^−^/^−^ thymocytes, leading to an antiproliferative response [15]. A later study by the same group proofed transgenic mice with elevated sIFNAR2 expression to be more susceptible to LPS-mediated septic shock, in which IFNβ plays a major role. Spleen cells of those transgenic mice overexpressing sIFNAR2a showed a faster, higher and more sustainable activation of STAT1 and STAT3, hence highlighting the agonistic abilities of sIFNAR2a [53]. Organ-dependent ratios of the mRNA for sIFNAR2a versus the mRNA for transmembranous IFNAR2 indicate an independent regulation from transmembranous IFNAR2 and a possibly organ specific biological function [15]. In humans, sIFNAR2a is reported to be elevated in systemic lupus [54] and various types of cancer [17] and to be associated with rather aggressive types of lung cancer [18]. Contradicting findings of elevated [16], decreased [55] and unchanged [54] sIFNAR2a levels in multiple sclerosis compared to healthy patients were reported. IFN therapy was further shown to induce elevated levels of sIFNAR2a compared to differentially treated multiple sclerosis patients and healthy donors [54]. This finding reflects what is seen during hepatitis C infection, in which sIFNAR2a expression was found to be predictive for the response to IFNα2 therapy [19]. One study reported a negative correlation between IFNAR2a expression and B cell exhaustion as well as impaired antibody production in the blood of HIV^+^ ART-treated, but not HIV^+^ ART naïve patients [56], which implicates ART rather than HIV itself as the primary effector and thus supports the data mentioned before [45]. Since we did not perform functional analysis of B cells in our samples, we cannot exclude the possibility of an association between sIFNAR2a levels and impaired B cell function. Given that we did not observe any difference between cART-naïve and cART-experienced patients, it is rather unlikely that HIV infection has any influence on the expression of the different IFNAR2 isoforms on LPMCs.

One factor influencing the diverse biological response of IFNα subtypes are their different affinities to IFNAR1 and IFNAR2, supported by the fact that affinity to IFNAR2 was reported to positively correlate with their antiviral activity against HIV [9]. To this end, we analyzed the effect of stimulating LPMCs with four IFNα subtypes, representing subtypes with high (IFNα14), middle (IFNα2 & IFNα8) and low (IFNα1) binding affinity to IFNAR2. Though IFNα8 and especially IFNα14 are highly antiviral against HIV, IFNα1 and IFNα2 exhibit only weak antiviral effects against HIV [9,31,33]. With all four tested subtypes, we did not see any differences in IFNAR2 expression on CD4^+^ T cells, CD8^+^ T cells, B cells or NK cells at 15 min, 30 min, 2 h and 24 h post stimulation (Fig. 4). Important factors for shaping the biological response of IFNα are the exposure time of the ligand, the receptor density expressed on the cell surface, the kinetics of its downregulation as well as the dose of the ligand [35]. Following IFN-stimulation with much higher concentrations than the reported EC_50_, we observed a downregulation of IFNAR2 on lymphocytes (unpublished data), however these concentrations exceed physiologically tolerable levels of IFNα, and thus are not applicable for therapeutic treatments. As recently reviewed [36], low doses of IFNα have only marginal effects on receptor expression, whereas high doses result in a stronger downregulation of receptor from the cell surface, since a considerably higher amount of receptor molecules is engaged [36] and thus showing a ligand induced dose dependent receptor downregulation.

With all IFNα subtypes possessing different affinities to both receptor subunits, it is likely that they also differ in their influence on ligand induced downregulation of the receptor. Several studies have shown downregulation of both IFNAR1 and IFNAR2 after ligand stimulation on cell lines [39] and on human PBMCs [44], however, only IFNα2 was tested in these studies. In contrast to our findings, Tochizawa *et al*. [44] observed decreased IFNAR2 expression on PBMCs following 2 h stimulation with 100 U/ml and 1000 U/ml IFNα2, the latter being equal to the EC_50_ for IFNα2 used in our experiments. As discussed above, it is possible that LPMCs react differently than PBMCs to IFNα stimulation and further elucidation of which cell types exactly exhibit the observed IFNAR2 downregulation is needed. Out of the four cell lines tested by Marijanovic *et al.* [39], IFNAR2 downregulation was only detected on Hek293T cells and on HeLa cells, but not on Daudi or Jurkat cells. Among those four, Daudi and Jurkat cells are the only ones originating from immune cells, with Daudi cells being Burkitt Lymphoma cells and Jurkat cells originate from human T cells. Thus, the expression profile of IFNAR1 and IFNAR2 on Daudi and Jurkat cells is more likely to resemble the expression profile of human *ex vivo* samples than the expression profile of HeLa and Hek293T cells. Additionally, the downregulation measured on HeLa and Hek293T cells was more pronounced for IFNAR1 than it was for IFNAR2. As mentioned previously, the expression of IFNAR1 on the cell surface is tightly regulated [47]. Even though we could not detect an effect of IFNα stimulation on the expression of IFNAR2, it is likely that IFNAR1 expression is more sensitive to IFNα subtype induced downregulation and its expression has greater influence in shaping the IFNα subtype dependent biological response.

In conclusion, neither HIV infection nor IFNα stimulation seem to influence the expression of IFNAR2, which therefore likely does not to affect the IFN responsiveness of cells in the gut during HIV infection.

## Acknowledgements

We would like to acknowledge Stefan Esser for providing plasma samples of HIV+ infected patients, the Westdeutsche Biobank for providing gut biopsies from patients at the University Hospital Essen and Barbara Bleekmann for excellent technical assistance. TZM-bl cells were a kind gift from the NIH AIDS reagent program.

